# Precise and Predictable DNA Fragment Editing Reveals Principles of Cas9-Mediated Nucleotide Insertion

**DOI:** 10.1101/135822

**Authors:** Jia Shou, Jinhuan Li, Qiang Wu

**Author notes:** Correspondence (Q.W.). Co-first author (Jia Shou, Jinhuan Li).

## Abstract

DNA fragment editing (DFE) or chromosomal engineering including inversions, deletions, and duplications by Cas9 with paired sgRNAs are important to investigate structural genome variations and developmental gene regulation, but little is known about the underlying mechanisms. Here we report that debilitating CtIP, which is thought to function in NHEJ, enhances precise DNA fragment deletion. By analyzing the inserted nucleotides at the junctions of DNA fragment inversions, deletions, and duplications, we find that Cas9 cleaves the noncomplementary strand with a flexible profile upstream of the PAM site and rationally-designed Cas9 nucleases have distinct cleavage profiles. Finally, Cas9-mediated nucleotide insertions of DFE are nonrandom and are equal to the combined sequences upstream of both PAM sites with predicted frequencies. Thus, precise and predictable DFEs could be achieved by perturbing DNA repair genes and using appropriate PAM configurations. These findings have important applications regarding 3D chromatin folding and enhancer insulation during gene regulation.

## Introduction

Organisms defend themselves against virus infection with adaptive immunity and maintain their genome integrity against genetic damage with DNA repair machinery. In bacteria and archaea, for example, clustered regularly interspaced short palindromic repeats (CRISPR) are a natural adaptive immune system against virus infection and phage conjugation (Deltcheva et al., 2011; Jiang et al., 2016; Jinek et al., 2012; Makarova et al., 2015; Marraffini and Sontheimer, 2010; Sternberg et al., 2015). For the type II system, a single endonuclease of the CRISPR associated protein (Cas9) is programed to cleave the invading viral genomes by crRNA (CRISPR RNA) and tracrRNA (trans-activating crRNA), generating blunt-ended double-strand breaks (DSBs) 3 base pairs upstream (-3 bp) of PAM (protospace adjacent motif, NGG for Cas9 from *Streptococcus pyogenes*) by the HNH and RuvC nuclease domains for the complementary and noncomplementary strands, respectively (Deltcheva et al., 2011; Gasiunas et al., 2012; Jinek et al., 2012; Makarova et al., 2015). In particular, the R-loop locking of HNH by guide RNA enables the cutting of the complementary strand at the exact -3 position upstream of PAM (Jiang et al., 2016; Sternberg et al., 2015).

In eukaryotes, Cas9 can be reprogramed for DNA-fragment editing (DFE) by paired sgRNAs to generate four DSB ends, which are repaired by several competing end-joining (EJ) pathways including canonical or classic non-homologous EJ (cNHEJ), alternative NHEJ (alt-NHEJ or MMEJ), and homologous recombination (Bétermier et al., 2014; Bhargava et al., 2017; Jasin and Haber, 2016), resulting in large DNA-fragment deletions and inversions as well as duplications (Bhargava et al., 2017; Canver et al., 2014; Huang and Wu, 2016; Kraft et al., 2015; Li et al., 2015a). This DFE method of chromosomal rearrangement or genome engineering is useful to probe 3D chromatin architecture and gene regulation; however, the underlying mechanisms are largely unknown (Dekker and Mirny, 2016; Franke et al., 2016; Guo et al., 2015; Huang and Wu, 2016; Lupiáñez et al., 2015).

Here we report that debilitating CtIP enhances precise DNA fragment deletion (DFD). In addition, in contrast to cleavage by HNH at the exact -3 position of the complementary strand (Jiang et al., 2016; Sternberg et al., 2015), we found that the cleavage by RuvC on the noncomplementary strand is flexible. Specifically, RuvC cleaves the noncomplementary strand at -3 as well as further upstream positions of PAM *in vivo*, resulting in blunt as well as non-blunt DSB ends with 5’ overhang. Through rational design of a set of engineered Cas9 nucleases, we achieved predictable DNA fragment inversion (DFI) *in vivo* based on the scissile profiles of RuvC. Finally, with this, we dissected the orientation and function of a composite enhancer containing two CBSs (CTCF-binding sites), revealing that one single CBS at topological domain boundary functions as an insulator to ensure specific long-distance chromatin-looping interactions between the distal enhancer and its target promoter. Thus, our mechanistic insights of Cas9 cleavage and of subsequent DSB-repair events should have broad implications for precise and predictable editing of millions of noncoding elements for 3D genome architecture and chromosomal rearrangement in genetic diseases.

## Results

### The Role of CtIP in Precise DFD Revealed by Screening DNA Repair Genes

DFD by Cas9 with paired sgRNAs has the advantage of avoiding single-sgRNA-guided Cas9 re-cutting (Paquet et al., 2016). Indeed, we observed many more precise junctions of DFD guided by paired sgRNAs (Guo et al., 2015; Li et al., 2015a). To further increase the efficiency of precise DFD, we screened a panel of seven DNA-repair genes and found that debilitating CtIP (C-terminal binding protein[CTBP]-interacting protein, also known as RBBP8: RB-binding protein 8, or Sae2) results in a significant increase of precise DFD (Figure 1). Specifically, we knocked out each of the seven DNA-repair genes and found, through next generation sequencing (NGS), that precise DFDs at the protocadherin (*Pcdh*) locus are significantly increased with CtIP debilitation (Figure 1A).

**Figure 1.**
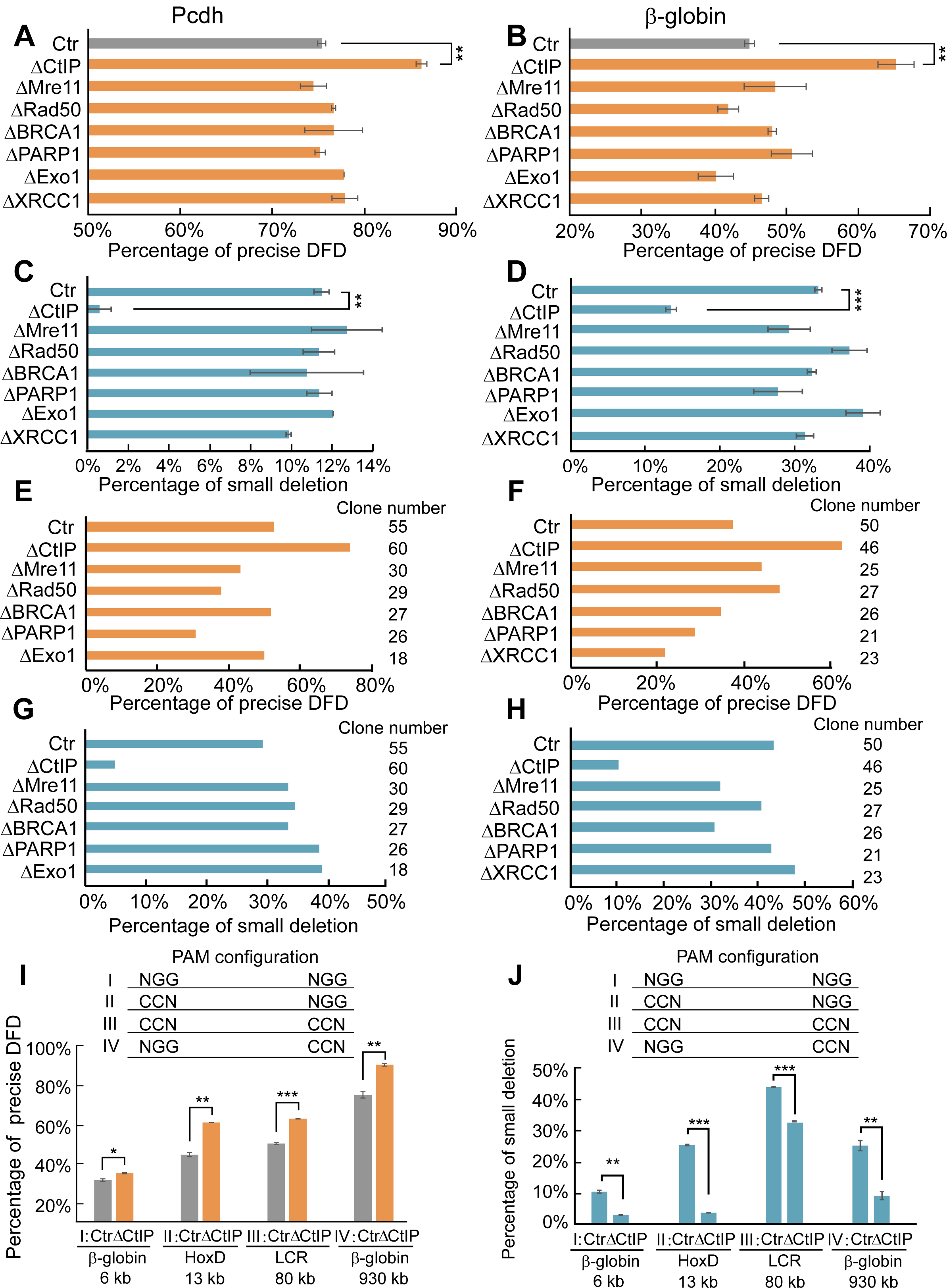
CtIP Debilitation Increases the Frequency of Precise DFD. (A-B) Quantitative analyses by NGS of the deletion junctions of a DNA fragment in the *Pcdh* or *β-globin* locus. (C-D) Quantitative analyses of small deletions at DFD junctions in the *Pcdh* and *β-globin* loci. (E-F) Confirmation of the increase of precise DFDs with CtIP debilitation by TA cloning and Sanger sequencing. (G-H) Confirmation of the decrease of small deletions at the DFD junctions with CtIP debilitation by TA cloning and Sanger sequencing. (I) CtIP deficiency leads to increased precise DFD of distinct DNA fragments of different sizes ranging from 6 kb to 930 kb by paired sgRNAs in the four PAM configurations. (J) CtIP debilitation leads to a significant decrease of small deletions at DFD junctions of distinct DNA fragments with different sizes ranging from 6 kb to 930 kb in all four PAM configurations. Data as mean ± SD, \**P* < 0.05, \*\**P* < 0.01, \*\*\**P* < 0.001; one-tailed Student’s *t*-test.

To see whether this is true for other loci, we screened the same panel of DNA-repair genes for precise DFD in the *β-globin* locus. Indeed, CtIP deficiency also enhances the efficiency of the *β-globin* precise DFD (Figure 1B). In addition, we found a corresponding decrease of small deletions and no significant alteration of insertions at the DFD junctions (Figures 1C-1D and S1A-S1B). We confirmed these NGS results by cloning and Sanger sequencing (Figures 1E-1H). Finally, CtIP debilitation results in no significant changes at the junctions of DNA fragment inversion or duplication (Figures S1C-S1H), suggesting that the mechanism of DFD is different from that of DNA-fragment inversion and duplication (*trans*-allelic translocation) (Ghezraoui et al., 2014).

We next investigated four combinatorial CRISPR PAM configurations with paired sgRNAs for distinct DNA fragments of different sizes in the *β-globin* and *HoxD* loci. We found that CtIP deficiency results in a significant increase of precise ligations and a corresponding decrease of small deletions at the DFD junctions, but no significant changes in the junctions of DNA fragment inversion and duplications in all four PAM configurations (Figures 1I-1J and S1I-S1L).

### Confirmation the role of CtIP in DFD by Triapine and CRISPR Single Cell Clones

To confirm the role of CtIP in DFD, we used triapine (3AP) to block CtIP activity and found that 3AP increases precise DFD and decreases the percentages of small deletions but no alteration of insertions for both the *Pcdh* and *β-globin* loci (Figures 2A-2D and S2A-S2B). However, there are no significant alterations in the junctions of DNA fragment inversion and duplication (Figures S2C-S2H).

**Figure 2.**
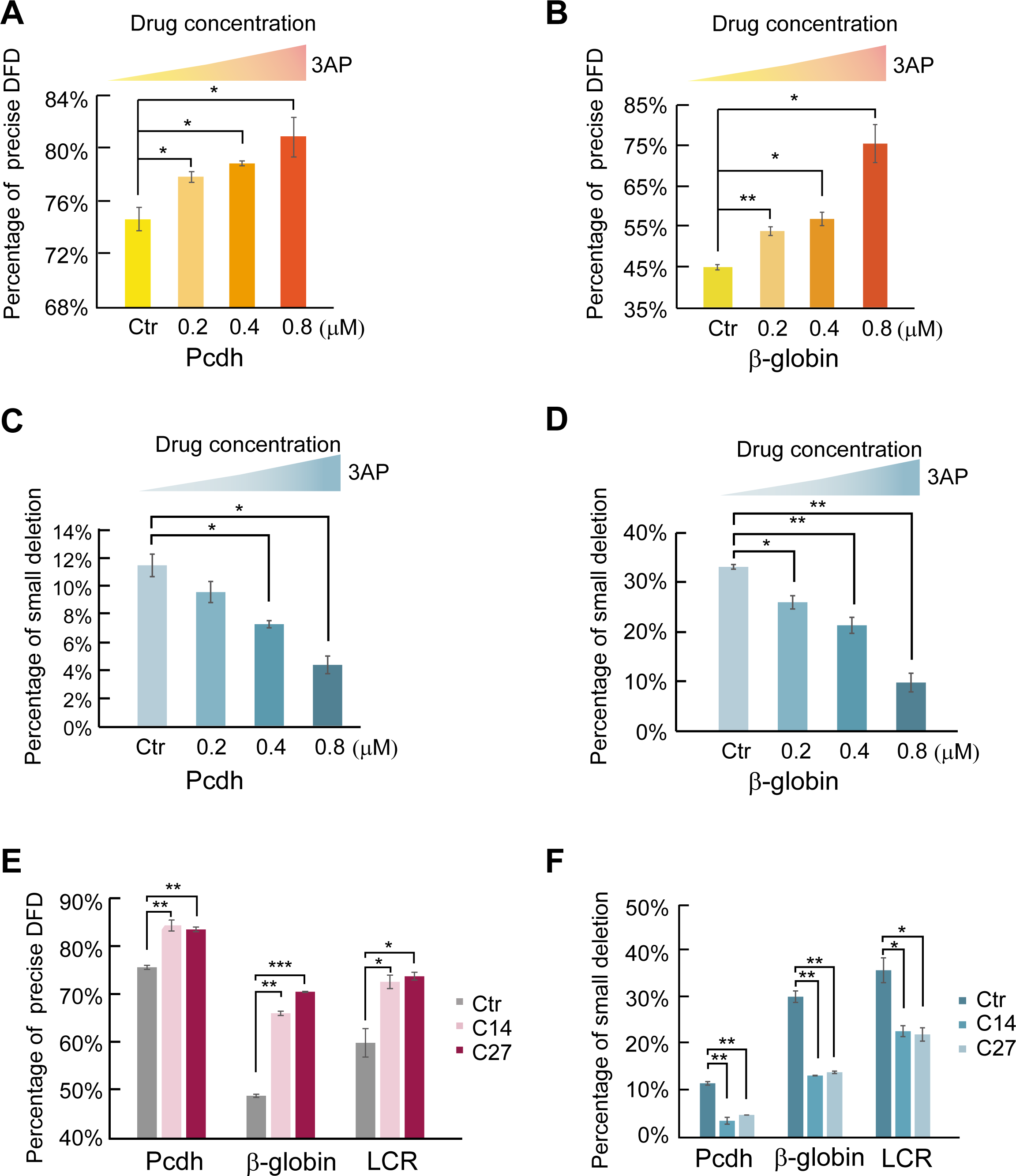
Confirmation of the Role of CtIP by Triapine and CRISPR Single Cell Clones. (A-B) Inhibition of CtIP activity by triapine (3AP), a drug that leads to inactivation of CtIP, increases the frequency of precise DFD in the *Pcdh* and *β-globin* loci. (C-D) CtIP inhibition by 3AP results in a significant decrease of small deletions at the DFD junctions in the *Pcdh* and *β-globin* loci. (E) CtIP-deficient cell clones (C14, C27) generated by CRISPR engineering display a significant increase of precise DFD frequency. (F) There is a significant decrease of small deletions in CtIP mutant cell clones at the *Pcdh*, *β-globin*, and its LCR. Data as mean ± SD, \**P* < 0.05, \*\**P* < 0.01, \*\*\**P* < 0.001; one-tailed Student’s *t*-test.

Finally, we screened for CtIP-mutant CRISPR cell clones and measured the precise DFD efficiency at *Pcdh*, *β-globin*, and its *LCR* (locus control region) by NGS. We obtained two CtIP-mutant CRISPR cell clones (C14 and C27) and found both of them display a significantly higher efficiency of precise DFD and a corresponding decrease of small deletions at DFD junctions (Figures 2E and 2F). However, these two cell clones display no significant alterations in the junctions of DNA fragment inversion and duplication (Figures S2I-S2N). Together, these data demonstrate that debilitating CtIP enhances precise DFD. Since DFD results from two competing DNA-repair pathways (cNHEJ and alt-NHEJ) (Bhargava et al., 2017) and CtIP facilitates or participates in strand resection in alt-NHEJ (Sartori et al., 2007), this suggests that cNHEJ is an error-free DSB repair pathway (Bétermier et al., 2014).

### Non-blunted end cleavage by Cas9

Taking advantage of the absence of Cas9 re-cutting in DFE (Paquet et al., 2016), in conjunction with NGS, we next investigated the EJ-repair junctional patterns of deletion, duplication, and inversion of a *Pcdh* DNA fragment (Figure 3). We first analyzed the pattern of indels at DFD junctions and found a striking pattern of nucleotide insertions (Figure 3A). Specifically, at the first position of insertions, there is a “G” nucleotide with a 99.11% frequency (Figure 3A). By contrast, there is a “T” with a frequency of only 0.89% at this position with no “C” or “A” at all. At the second position of insertions, there is 100% “G” with no insertion of “T”, “C”, or “A” (Figure 3A). Remarkably, there is an “A”, “C”, “T”, “G”, or “C” nucleotide each with a 100% frequency at the 3^rd^, 4^th^, 5^th^, 6^th^, or 7^th^ position of insertions, respectively, with no other nucleotides at all (Figure 3A). Thus, the nucleotide insertions at each position are nonrandom and may reflect the cleavage pattern of RuvC on the noncomplementary strand. Indeed, the actual inserted nucleotides of “+G”, “+GG”, “+AGG”, “+CAGG”, “+TCAGG”, “+GTCAGG”, and “+CGTCAGG” all match perfectly with the nucleotide sequences of the noncomplementary strand at the -4, -5, -6, -7, -8, -9, and -10 positions upstream of the second PAM site, respectively (Figure 3A). These observations strongly suggest that the inserted nucleotides at the DFD junctions result from the flexible cleavage of RuvC of the noncomplementary strand at the -4, -5, -6, -7, -8, -9, and -10 positions.

**Figure 3.**
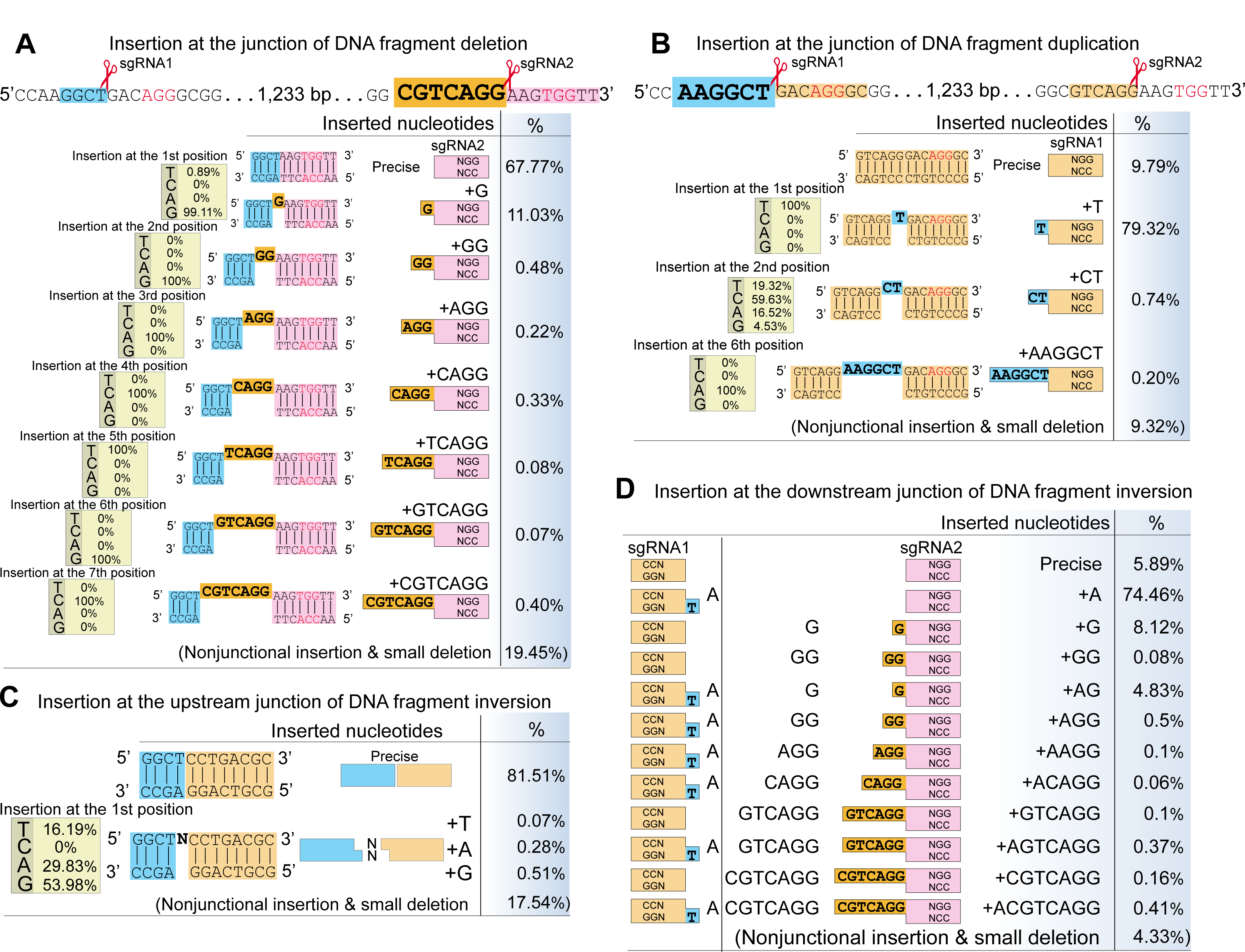
Non-random Nucleotide Insertions at DFE Junctions *in vivo*. (A) The inserted nucleotides at DFD junctions are resulted from the cleavage by RuvC of the noncomplementary strand upstream of the **second** PAM site. The sequences of a targeted *Pcdh* DNA fragment (1,233 bp) are shown with PAM in red. The inserted nucleotides detected by NGS are highlighted in the yellow background and are shown with their frequencies. The relative ratios of “T”, “C”, “A”, “G” at each position of insertions are also shown. (B) The inserted nucleotides at the junctions of DNA fragment duplications are from the cleavage by RuvC of the noncomplementary strand upstream of the **first** PAM site. The inserted nucleotides are highlighted in the blue background and are shown with their frequencies. (C) Nonspecific insertions at the upstream junctions of DFI. (D) The inserted nucleotides at the downstream junctions of DFI are shown with their frequencies. They are resulted from the combined insertions at junctions of both DNA-fragment deletion (Cas9 with sgRNA2) and duplication (Cas9 with sgRNA1).

We then analyzed nucleotide insertions at the junctions of DNA-fragment duplications and found a pattern similar to that of DFD. Not only is there a striking nucleotide bias of “T”, “C”, or “A” with a frequency of 100%, 59.63%, or 100% at the 1^st^, 2^nd^, or 6^th^ position of insertions, respectively; but the inserted nucleotides of “+T”, “+CT”, and “+AAGGCT” also match perfectly with the nucleotide sequences of the noncomplementary strand upstream of the first PAM site (Figure 3B). We concluded that the inserted nucleotides at the junctions of DNA-fragment duplication result from the flexible cleavage of RuvC of the noncomplementary strand at the -4, -5, or -9 position upstream of the first PAM site. Although there is no specific nucleotide-insertion pattern at the upstream DFI junctions, we found a remarkable pattern of nucleotide insertions at the downstream DFI junctions (Figures 3C and 3D). Namely, the inserted nucleotides match the combined sequences of the nucleotide insertions at junctions of both DNA-fragment deletion and duplication (Figure 3D).

Finally, we edited four additional DNA fragments of different sizes ranging from ∼700 bp to ∼930 kb in distinct loci and found remarkably-similar cleavage patterns (Figures S3 and S4). Taken together, these data strongly suggest that, in addition to cleavage of the noncomplementary strand at the -3 position upstream of the PAM site, RuvC also cleaves of the noncomplementary strand at positions further upstream *in vivo*.

### Altered Cleavage Profiles of Engineered Cas9 Nucleases by Rational Design

To confirm the non-blunted cutting pattern of Cas9 *in vivo*, we used a structure-based rational design to alter the cleavage pattern of the RuvC domain. Recent structural and molecular studies revealed that allosteric conformational changes between RuvC and HNH for R-loop formation are essential for Cas9 catalytic activation and that the two linker regions between these domains are mostly disordered or helical (Anders et al., 2014; Jiang et al., 2016; Jinek et al., 2014; Nishimasu et al., 2014; Sternberg et al., 2015). We reasoned that, by perturbing these two linker regions and carefully preserving the catalytic activity of RuvC and HNH, we may be able to design an engineered Cas9 with altered scissile patterns of RuvC on the noncomplementary strand.

We screened twenty engineered Cas9 nucleases with mutations of residues in the two linker regions and found seven of them with altered scissile profiles for each of the ten sgRNAs we tested (Figures 4A and 4B). We then calculated the scissile profile for each engineered nuclease with these ten sgRNAs and found that each has a distinct scissile profile (Figure 4C). Finally, we tested these engineered nucleases in our CtIP CRISPR cell clone and found that CtIP deficiency results in a significant increase of the percentage of precise DFD and a concomitant decrease of small deletions for each rationally-designed Cas9 nucleases (Figures 4D-4G). These data suggest that the cleavage profile of Cas9 could be altered by inducing a conformational change through engineering the linker regions between the HNH and RuvC domains.

**Figure 4.**
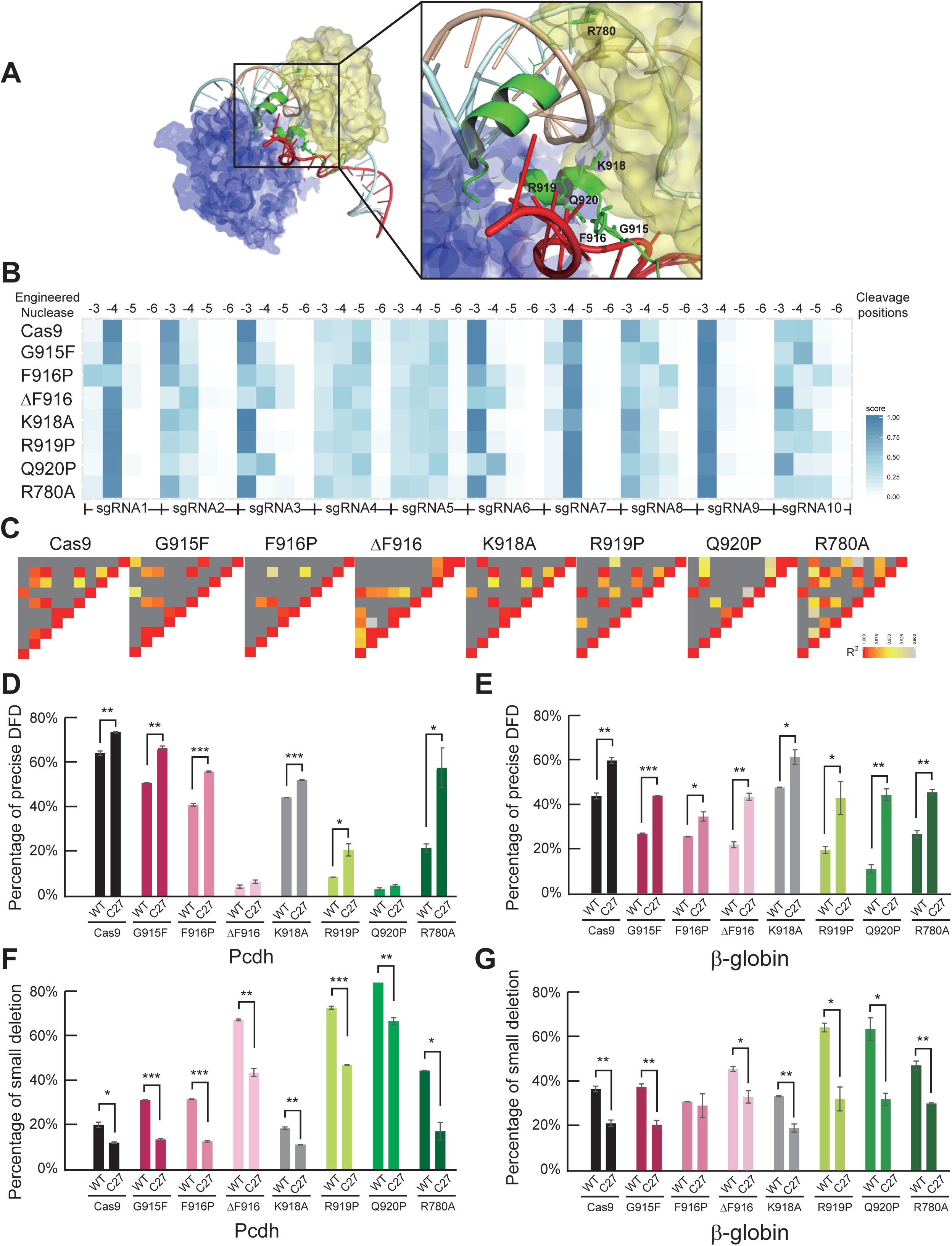
Altered Scissile Profiles of Rationally Engineered Cas9 Nucleases. (A) A surface representation of the crystal structure of SpCas9 (PDB 5F9R) (Jiang et al., 2016) showing the two linker regions (depicted in green ribbon) between the HNH (yellow) and RuvC (blue) domains. The noncomplementary strand is depicted in red, the complementary strand in cyan, and sgRNA in wheat. (B) Altered scissile profiles at -3 to -6 positions on the noncomplementary strand of engineered Cas9 nucleases. (C). Each engineered Cas9 nuclease display a unique scissile profiles for the ten sgRNAs tested. (D-G) The engineered Cas9 nucleases display a significant increase of precise DFD and a concomitant decrease of small deletions in the *Pcdh* and *β-globin* loci in the *CtIP*-deficient cell clone. Data as mean ± SD, \**P* < 0.05, \*\**P* < 0.01, \*\*\**P* < 0.001; one-tailed Student’s *t*-test.

### Predictable DFE

Deep sequencing of duplication junctions revealed that all of them except K918A result in a decreased frequency of cleavage of the noncomplementary strand at the -3 position (precise) and that four of them have an increased frequency of cleavage at the -4 position (insertion of “+C”) upstream of the first PAM site (Figures 5A and 5B). In addition, most of them also have an increased frequency of cleavage at the -5 position (insertion of “+GC”) (Figure 5B).

**Figure 5.**
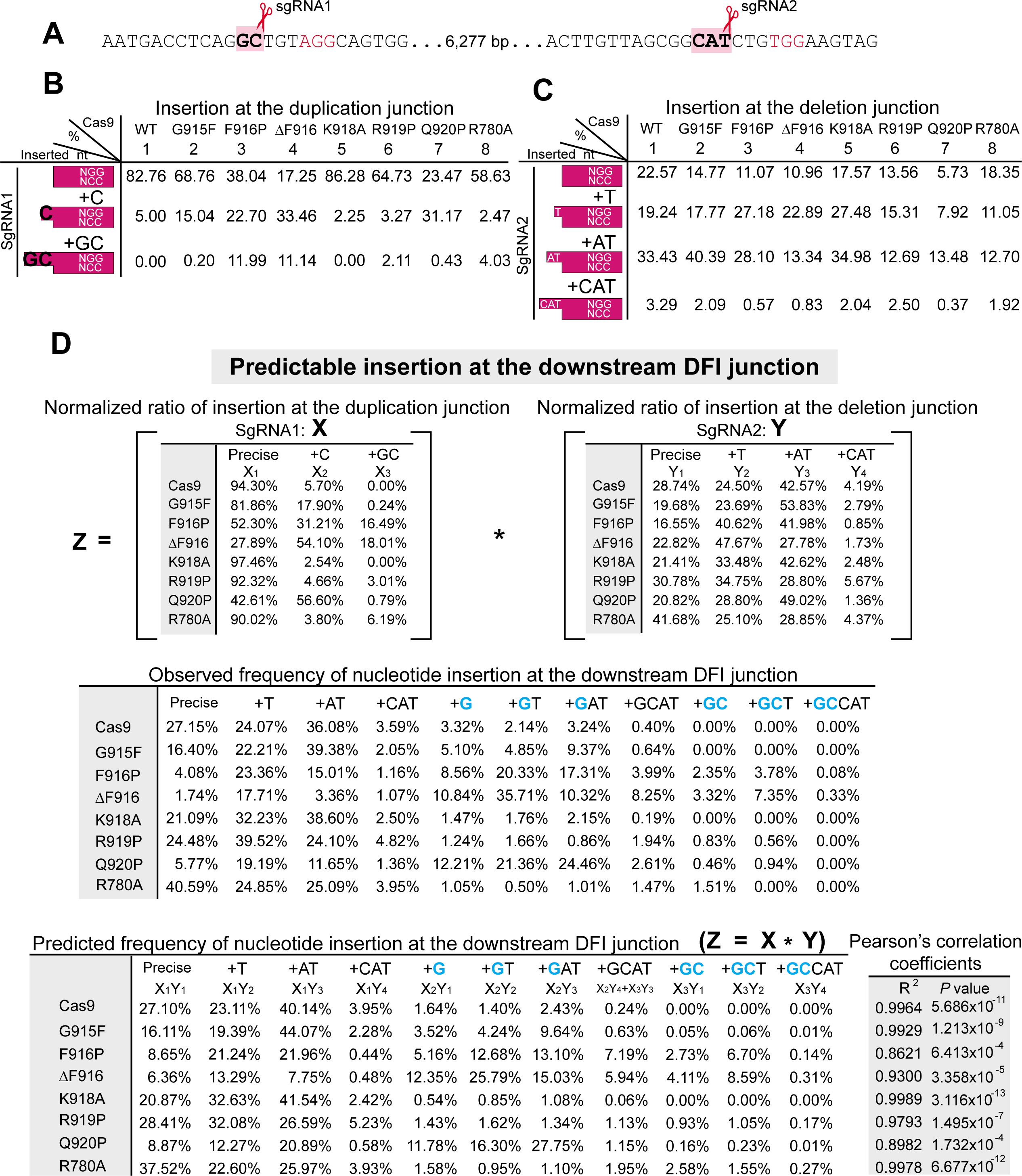
Predictable Insertions at the Downstream DFI Junctions. (A) The sequences of a targeted *β-globin* DNA fragment (6,277 bp) are shown with PAM in red. (B-C) Shown are the inserted nucleotides and their frequencies at the duplication or deletion junctions of engineered Cas9 nucleases. (D) Predictable Cas9-mediated nucleotide insertions at the downstream DFI junctions. The observed frequencies of the downstream insertions at the DFI junction are predicted to be equal to the product of multiplication of the normalized insertion frequencies at the both duplication and deletion junctions. Note that the nucleotide insertion of “+GCAT” are from both “X_2_Y_4_” (the combined insertion of “+**G**” and “+CAT”) and “X_3_Y_3_” (the combined insertion of “+**GC**” and “+AT”). The Pearson’s product-moment correlation coefficients and their *P* values for all of the engineered Cas9 nucleases are indicated on the right.

Deep sequencing of deletion junctions revealed that all of the seven engineered Cas9 nucleases have a decreased frequency of cleavage at the -3 position and the scissile profiles at -4, -5, and -6 positions upstream of the second PAM site are also altered (Figure 5C).

Remarkably, nucleotide insertions at the downstream junctions of DFI perfectly match the combined insertions at the junctions of both DNA-fragment deletion and duplication with the expected frequency (Figure 5D). These observations further support that Cas9 generates both blunt and non-blunt DSB ends with a 5’ overhang *in vivo*. Indeed, we could predict the frequency of inserted nucleotides at downstream inversion junctions for all of the seven engineered Cas9 nucleases (Figure 5D).

Finally, we edited two additional DNA fragments of different sizes ranging from ∼700 bp to ∼930 kb and found the same remarkable principle of Cas9-mediated nucleotide insertions (Figures S5 and S6). Taken together, we conclude that precise and predictable Cas9-mediated DFEs can be achieved by engineered Cas9 with paired sgRNAs.

### Single CBS sites at Topological Domain Boundaries Determine Chromatin Looping Directions

This precise and predictable DFE method allows a detailed investigation of the role of enhancer and insulator orientation in 3D genome architecture and gene regulation. The location and relative orientation of CBSs genome-wide determine the 3D chromosomal architecture (de Wit et al., 2015; Guo et al., 2015; Sanborn et al., 2015). The human *Pcdh* locus comprise 53 tandem genes organized into three sequentially-linked clusters (*Pcdhα*, *β*, and *γ*) (Wu and Maniatis, 1999) and two (*Pcdhα* and *Pcdhβγ*) CTCF/cohesin-mediated chromatin contact domains (CCDs or subTADs) (Guo et al., 2015). These clustered *Pcdh* genes are crucial for dendritic self-avoidance and tiling during brain development (Lefebvre et al., 2012). The orientation and location of a repertoire of *Pcdh* CBSs determine the directional looping between the distal *HS5-1* enhancer and its target promoters in the *Pcdhα* variable region (Guo et al., 2015). This composite enhancer comprises two tandem CBSs (*HS5-1a* and *HS5-1b*) both in reverse orientation and numerous TF binding sites that are clustered in a 1-kb region (Figure 6A) (Guo et al., 2015).

**Figure 6.**
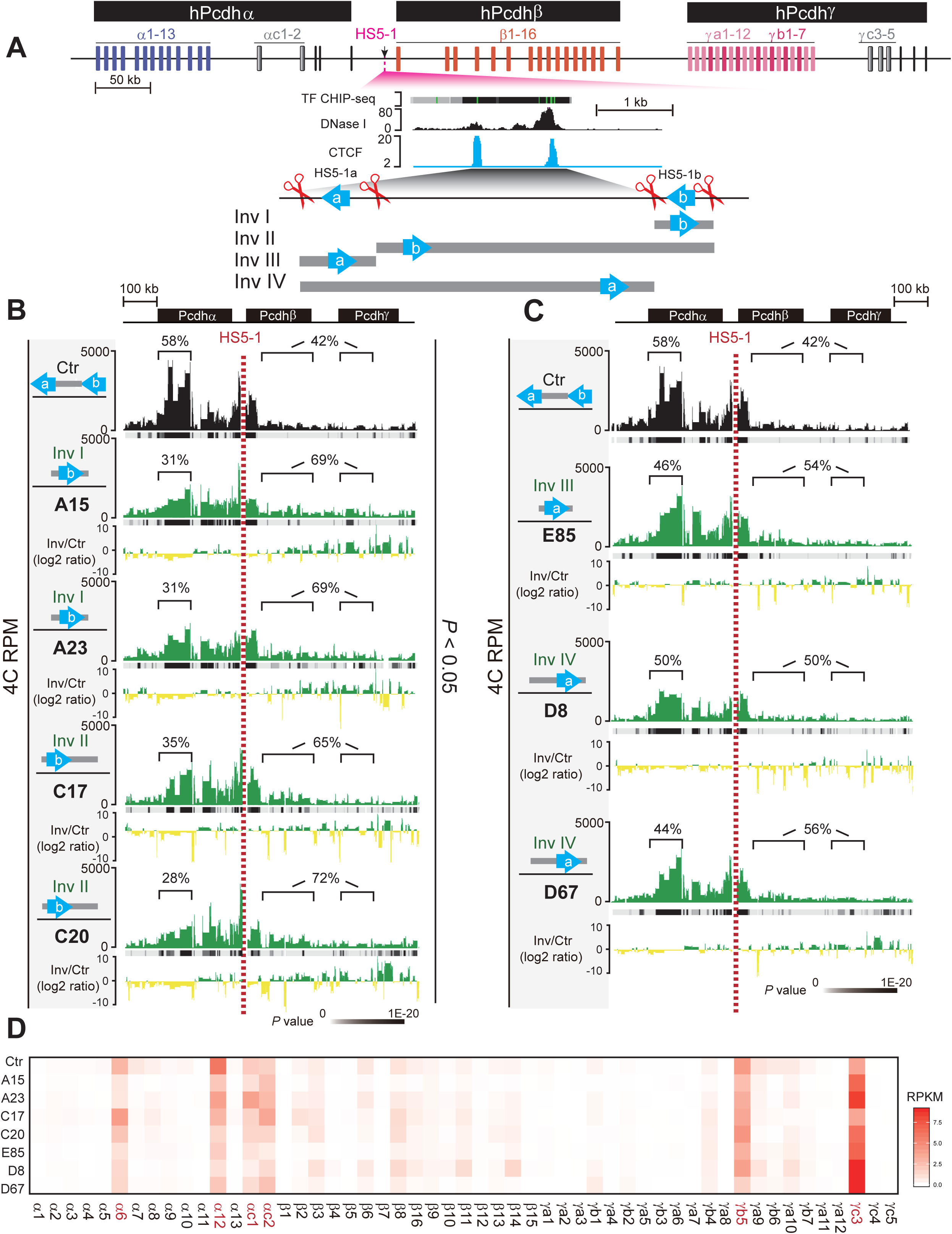
Dissection of the *Pcdh HS5-1* Enhancer Architecture by DFI Reveals the Insulational Role of the Single Boundary CTCF Site. (A) Schematics of the three human *Pcdh* gene clusters. The composite *HS5-1* enhancer and its various CRISPR inversions are enlarged. Two tandem CBSs in a reverse orientation (*HS5-1a* and *HS5-1b*) are indicated. (B) Shown are chromatin-interaction profiles in the wild-type control (Ctr) and CRISPR cell lines with inversion of the CBS *HS5-1b* alone (Inv I) or the *HS5-1b* combined with the middle region (Inv II) using the *HS5-1* enhancer as an anchor. The significance of interactions (*P* value) is shown under the reads density. The log2 ratios between each inversion cell clone and wild-type cells are also indicated. (C) Shown are chromatin-interaction profiles in wild-type (Ctr) and CRISPR cell lines with inversion of the CBS *HS5-1a* alone (Inv III) or the *HS5-1a* combined with the middle region (Inv IV) using the *HS5-1* enhancer as an anchor. (D) Heatmap shows the alteration of expression patterns of members of the *Pcdh* gene clusters resulting from CRISPR inversion of different enhancer elements by RNA-seq experiments.

To date, there is no evidence that the relative orientation of single CBS sites can determine chromatin-looping directions between enhancers and promoters (de Wit et al., 2015; Guo et al., 2015; Sanborn et al., 2015). To this end, we designed a series of paired sgRNAs to dissect the *HS5-1* enhancer architecture (Figure 6A). We screened single-cell CRISPR clones for inversion of the single CBS *HS5-1a* or *HS5-1b* site as well as its combination with the middle region of the *HS5-1* enhancer. Remarkably, inversion of *HS5-1b* alone or with the middle region results in a significant decrease of DNA-looping interactions between the *HS5-1* enhancer and the *Pcdhα* promoters, and a corresponding increase of DNA-looping interactions with the *Pcdhβγ* promoters (Figure 6B). However, inversion of *HS5-1a* alone or combined with the middle region results in no significant alteration of directional DNA-looping interactions with either *Pcdhα* or *Pcdhβγ* promoters (Figure 6C). Thus, the relative orientation of the single CBS *HS5-1b* at the *Pcdhα* CCD or subTAD boundary determines the chromatin-looping directions between the distal enhancer and its target promoters.

To assess the functional role of the altered chromatin looping, we performed the RNA-seq experiments and found that each CRISPR DFI cell line with disrupted architecture of the *HS5-1* enhancer displays altered patterns of *Pcdh* gene expression, consistent with previous studies that subtle structural changes could result in alternations of gene expression pattern (Figure 6D) (de Laat and Duboule, 2013). Finally, a series of CRISPR inversions of progressive numbers of CBSs in the *β-globin* locus demonstrate that only those inversions covering the CBS at a TAD boundary (CBS15) switch chromatin-looping directions (Figure S7). We conclude that single CBS sites at either TAD or SubTAD boundaries function as insulators.

## Discussion

Chromosomal engineering or large genomic fragment editing, including deletions, inversions, duplications, and translocations, could be achieved by using the conventional Cre-LoxP system with low efficiencies (Wu et al., 2007; Zheng et al., 2000). Recent development in genome editing by ZFN, TALEN, and CRISPR enabled rapid technical advance for chromosomal rearrangement or segmental editing (Blasco et al., 2014; Canver et al., 2014; Choi and Meyerson, 2014; Cong et al., 2013; Gupta et al., 2013; Kraft et al., 2015; Lee et al., 2012; Li et al., 2015a; Park et al., 2016; Torres et al., 2014; Xiao et al., 2013; Yang et al., 2013). In particular, DFE has been used to investigate developmental gene regulation and to model human genetic diseases (Franke et al., 2016; Guo et al., 2015; Lupiáñez et al., 2015; Maddalo et al., 2014; Park et al., 2015; Tai et al., 2016); however, the underlying mechanism for CRISPR DFE remains unknown and the nucleotide insertions at the junctions are assumed to be random. We found here that CtIP deficiency increases the frequency of precise DFD by Cas9 with paired sgRNAs *in vivo*. In addition, we found that CRISPR/Cas9-mediated nucleotide insertions are nonrandom and can be predicted based on the scissile profiles of RuvC on the noncomplementary strand. Our rational design and *in vivo* cleavage data are consistent with the allosteric conformation change between the HNH and RuvC domains for catalytic activation of Cas9 nucleases (Jiang et al., 2016; Sternberg et al., 2015). Finally, we found, through dissection of a composite enhancer, that single CBS sites at topological boundaries function as an insulator.

Based on our findings that RuvC cleaves the noncomplementary strand further upstream of PAM (Figures 3-4 and S3-S4) and that Cas9-mediated nucleotide insertions at the downstream DFI or duplication junctions are equal to the combined sequences with expected frequencies upstream of both PAM sites for the PAM configurations of NGG-NGG or NGG-CCN, respectively (Figures 5 and S5-S6); we propose that Cas9-mediated nucleotide insertions could be achieved for the other two PAM combinations of CCN-NGG and CCN-CCN (Figure 7). In addition, these inserted nucleotide sequences are generated by filling in of nucleotides through 5’-3’ extension by unknown DNA polymerases and subsequent precise ligation by DNA ligases during DNA DSB repair (Figure 7). Through these mechanisms, we could achieve predictable DFE for all four PAM configurations.

**Figure 7.**
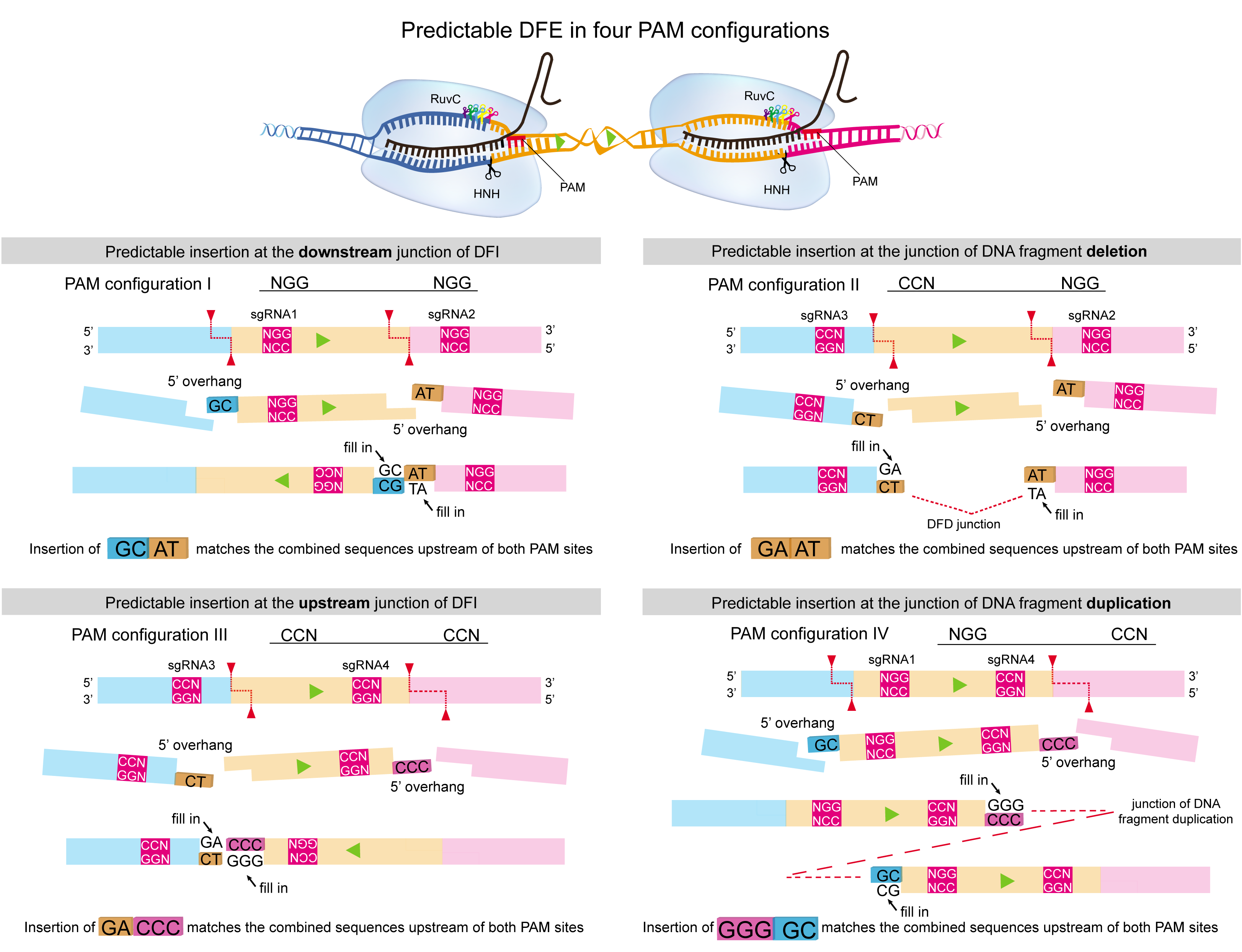
A Model of Predictable Cas9-mediated Nucleotide Insertions for the Four PAM Configurations. Shown are schematics of predictable DFEs by Cas9 with paired sgRNAs in four PAM configurations. Through filling in of DSB ends by an unknown polymerase and ligation, Cas9-mediated nucleotide insertions match the combined sequences upstream of both PAM sites for all four PAM configurations.

Our data suggest that, consistent with previous *in vitro* evidence (Jinek et al., 2012; Li et al., 2015b), Cas9 cleaves the dsDNA target to generate blunt as well as previously unrealized non-blunt ends with 5’ overhangs *in vivo*, revealing a striking similarity of staggered cutting patterns by the evolutionarily-related class 2 CRISPR effector proteins of Cas9, Cpf1, and C2c1 (Makarova et al., 2015; Yang et al., 2016; Zetsche et al., 2015). Furthermore, predictable insertions at DFI junctions based on the scissile profiles of RuvC of engineered nucleases on the noncomplementary strand reveal a general principle of Cas9-mediated nucleotide insertions. These findings on Cas9 cleavage and DSB-repair outcomes lay a foundation to improve DFE for probing 3D genome architecture and gene regulation as well as for investigating mechanisms of DNA damage response pathways.

## METHODS

### Cell culture

HEK293T cells were cultured in Dulbecco’s modified Eagle’s medium (HyClone) containing 10% FBS (Gibco) and 1% penicillin– streptomycin (Gibco). HEC-1-B cells were cultured in Eagle’s Minimum Essential Medium (Gibco) containing 10% FBS (Gibco), 1% penicillin– streptomycin (Gibco), 2 mM glutamine (Gibco), and 1 mM sodium pyruvate (Sigma). All cells were cultured at 37°C and 5% (v/v) CO_2_. Cells were plated at a density of approximately 4 × 10^5^ cells per well in 12-well plates and cultured for 24 hours for transfection.

### DNA-repair genes screening for precise DFD

First, plasmids for paired sgRNAs targeting the exons of each repair gene of *CtIP*, *Mre11*, *Rad50*, *BRCA1*, *PARP1*, *Exo1*, and *XRCC1* were designed (Supplemental Table S1) and constructed as described previously (Li et al., 2015a). In addition, a pair of sgRNAs targeting specific DNA fragments of the *Pcdh* as well as the *β-globin* loci (Supplemental Table S1) were also generated as described previously (Li et al., 2015a). HEK293T cells were transfected with plasmids encoding SpCas9 (0.8 μg), two sgRNAs targeting specific DNA fragment (each 0.6 μg) and two sgRNAs targeting the exons of each repair gene (each 0.6 μg) by Lipofectamine 2000 (Life Technology) in 12-well plates. The transfected cells were cultured for 48 hours and then harvested. Genomic DNA was extracted by Genomic DNA Purification kit (Promega) for analyses.

### NGS for DFD junctions

The *Pcdh* and *β-globin* DFD junctions were amplified with the high-fidelity DNA polymerase (KOD-Plus-Neo, TOYOBO) by specific primer pairs with P5 and P7 Illumina adapters (Supplemental Table S1). PCR products were purified by the High-Pure Purification Kit (Roche). Libraries were barcoded and sequenced on the Hiseq X-Ten platform. Reads of each sample were de-multiplexed and adaptor sequences were clipped. Reads were mapped to reference sequences of precise DFD junctions (Precise junctions are defined as direct ligations between two blunt DSB ends generated by Cas9 cutting at -3 bp upstream of the PAM site). Indels (insertions and deletions) of junctions were called by the Varscan2 program (Koboldt et al., 2012). The percentages of precise DFD were calculated. A total of 328 Illumina libraries were sequenced for analyzing the role of CtIP in precise DFD.

### TA cloning

TA cloning was used to confirm the results of NGS. Briefly, PCR products with a single adenosine attached to the 3′-ends were amplified by the Taq DNA polymerase and directly ligated into pGEM-T easy vector (Promega) by T4 DNA ligase. The ligated products were transformed with Stbl3 (F^-^ *mcr*B*mrrhsd*S20(rB^-^,mB^-^)*rec*A13*sup*E44*ara*-14*gal*K2*lac*Y1*pro*A2*rps*L20(Str^R^) *xyl*-5λ*leumtl*-1) competent cells. About 30 clones for each DNA-repair gene were analyzed by Sanger sequencing (Figure 1). The results of TA-cloning were consistent with NGS. Thus, all subsequent experiments were analyzed by NGS.

### PAM configuration

A pair of plasmids encoding paired sgRNAs for each PAM configuration targeting *HoxD* or *β-globin* DNA fragments (Supplemental Table S1) were generated as described previously (Li et al., 2015a). After transfection by Lipofectamine 2000 (Life Technology), genomic DNA was extracted and analyzed by NGS for precise DFD.

### Drug inhibition of the CtIP activity

Triapine (3-aminopyridine-2-carboxaldehyde thiosemicarbazone, 3AP) is a small-molecule inhibitor that was used to block CtIP phosphorylation (Lin et al., 2014). HEK293T cells were treated with 3AP at the final concentrations of 0.2 μM, 0.4 μM, and 0.8 μM at the time of plasmid transfection and continued culturing for 24 hours. The DFD junctions were then analyzed by NGS.

### Single cell screening of *CtIP* mutant by CRISPR

Generation of CRISPR single-cell clones of *CtIP* mutants were as previously described (Guo et al., 2015; Li et al., 2015a). Briefly, HEK293T cells were transfected with Cas9 and two sgRNAs targeting the exons of *CtIP*. After 2 day of transfection, puromycin (2 mg/ml) was added and cells were continued to culture for additional 4 days. The cells were then maintained in the puromycin-free medium for another 8 days and harvested to seed in 96-well plates at an approximately density of one cell per well. The single cells were cultured for another 10 days to grow to single-cell colonies. Single-cell clones were then screened to obtain the *CtIP* mutant cell lines. Of 96 single-cell clones picked, two CtIP mutant clones (C14 and C27) were identified. Precise DFD was then tested in these CtIP mutant clones.

### Rational design of Cas9 nuclease

We reasoned that mutating the two linker regions of Cas9 between HNH and RuvC catalytic domains while keeping their catalytic center unchanged may result in alteration of the scissile profile. Engineered Cas9 nucleases with substitution or deletion of specific residues within the two linker regions were rationally designed based on the crystal structures of SpCas9 (PDB accession number: 5F9R). Briefly, primer pairs for cloning engineered Cas9 nucleases (Supplemental Table 1) were designed by either substituting or deleting the codon for a specific residue in the two linker regions. Wild-type SpCas9 plasmids were used as templates to amplify mutant DNA fragments with the Q5 polymerase (NEB). The SpCas9 templates were then digested by DpnI. The amplified PCR products were ligated with T4 DNA ligase (NEB) at room temperature for 5 minutes and were transformed with the Stbl3 competent cells. In total we designed 20 Cas9 mutants and found 7 of them with altered scissile profiles. All cloned plasmids encoding engineered Cas9 nucleases were confirmed by Sanger sequencing.

### Single-cell screening of DNA fragment inversions by CRISPR

CRISPR single-cell inversion clones were screened as previously published (Guo et al., 2015; Li et al., 2015a). For the *Pcdh* locus, we obtained 2 (A15 and A23), 1 (E85), 2 (C17 and C20), and 2 (D8 and D67) inversion HEC-1-B clones of CBS *HS5-1b* and *HS5-1a* as well as their combination with the enhancer middle regions from 44, 85, 67, and 77 single-cell clones, respectively. For the *β-globin* locus, we obtained 3 (A3, A29 and A49), 1 (B36), 2 (E19 and E37) inversion HEK293T clones of CBS15, CBS14-15, and CBS13-14 from 49, 40, and 40 single-cell clones, respectively.

### Circularized chromosome conformation capture (4C)

The libraries for 4C-seq were generated as described previously (Guo et al., 2012; Guo et al., 2015; Hagège et al., 2007; Splinter et al., 2012). Briefly, a total of 10^7^ cells of HEK293T cells or HEC-1-B cells were cross-linked by formaldehyde and lysed to expose cell nucleus. The cross-linked nuclei were digested with HindIII or EcoRI while rotating at 900 rpm overnight and then ligated with T4 DNA ligase (NEB). After reverse cross-linking, the purified genomic DNA was digested with a 4-bp cutter (DpnII or NlaIII) and were ligated again. The re-ligated products were purified by the High Pure DNA Purification kit (Roche). Finally, 4C-seq libraries were amplified by inverse PCR using the high-fidelity DNA polymerase by a pair of primers each with a P5 or P7 Illumina adapter (Supplemental Table S1). The amplified libraries were sequenced on a Hiseq X-Ten platform. Reads were mapped to the human genome (hg19) using the Bowtie2 program (Langmead and Salzberg, 2012). The r3Cseq program in the R/Bioconductor package (Thongjuea et al., 2013) was used to analyze the long-range chromatin-looping interactions. All 4C-seq experiments were performed with replications.

### RNA-seq

RNA-seq experiments were performed as previously described (Guo et al., 2015). Briefly, about 4 × 10^5^ cells were used for each RNA-seq. Total RNA were prepared using an RNeasy kit (Qiagen). Polyadenylated mRNAs were then selected by the oligo dT beads (NEB) for library construction. After reverse transcription of polyA mRNAs with a polyT primer, the cDNA libraries were barcoded and sequenced on the Illumina Hiseq X ten platform. Specific index primers (Supplemental Table S1) were used to multiplex RNA-seq samples. Sequenced reads were align to the human genome (GRCh37) using the TopHat software (v2.0.14) (Trapnell et al., 2009) with the parameters of -N 0 –g 1 –x 1. The expression levels of each transcript were calculated using the Cufflinks software (v2.2.1) (Trapnell et al., 2010) with the default parameters. The heatmap was generated using the R program to display expression levels of members of the *Pcdh* gene clusters.

### NGS and statistical analyses

All experiments for NGS were performed with biological replicates. In total, 1024 libraries were constructed and sequenced by NGS for analyzing of DFE junctional patterns. For CtIP experiments, *P-* values were calculated using the one-tailed Student’s *t*-test. For predictable DFE junctions, we assume that the probability of ligations of any two DSB ends by DNA-repair machineries are equal. Thus, the ratio of the DNA repair events for DFE junctions is equal to the product of the percentages of two DSB ends to be ligated. Pearson’s product-moment correlation coefficients (PPMCC) was used to compare sets of nucleotide-insertion frequencies of engineered Cas9 nucleases from NGS with corresponding sets of predictions that are equal to the product of multiplication of the normalized insertion frequencies. The correlation coefficients and *P* values were generated by the R program.

### Cas9 structural analysis

The structural representation in Figure 4A was generated by PyMOL (http://www.pymol.org/). The coordinates of the structure of SpCas9 with sgRNA and dsDNA is from PDB accession number 5F9R (Jiang et al., 2016). We removed the recognition domain and PAM-interaction domain for clarity. The HNH and RuvC nuclease domains are shown in surface representation. The two linker regions are shown in ribbon. The side chains of mutated or deleted residues are shown and labelled with single-letter amino acid code.

## Author Contributions

Q.W. conceived the experimental design and wrote the manuscript with the help of S.J and J.L., who also performed all the experiments.

## Acknowledgments

We thank Profs Dan Czajkowsky and Yanli Wang for critical reading of the manuscript and all members of the Wu lab for discussion. This work was supported by grants from NSFC (31630039, 91640118, and 31470820).

---

**Figure S1.**
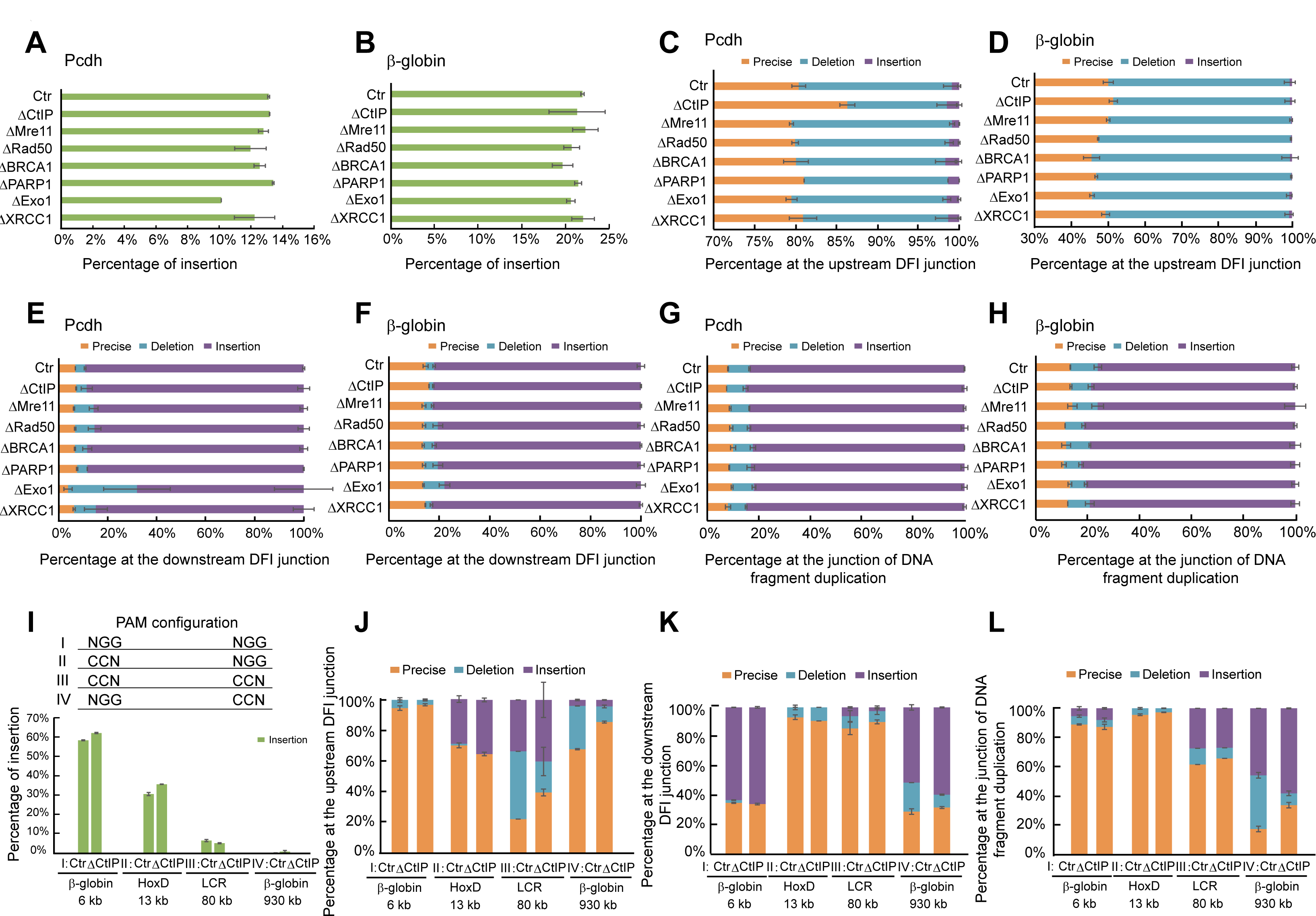
Characterization of Junctions of DNA Fragment Deletion Inversion, and Duplication with Inactivation of a Set of DNA-repair Genes and CtIP Debilitation in Four PAM Configurations, Related to Figure 1. (A-B) Quantitative analyses of insertions by NGS at DFD junctions in the *Pcdh* and *β-globin* loci. (C-D) Analyses of the frequency of precise ligations and indels by NGS at the upstream junctions of DFI in the *Pcdh* and *β-globin* loci. (E-F) Analyses of the frequency of precise ligations and indels by NGS at the downstream junctions of DFI. (G-H) Analyses of the frequency of precise ligations and indels by NGS at the junctions of DNA fragment duplication. (I) CtIP debilitation results in no significant alteration of insertions at DFD junctions. (J-L) CtIP debilitation results in no significant alteration of precise ligations and indels at the upstream and downstream DFI junctions as well as DNA-fragment-duplication junctions. Data as mean ± SD.

**Figure S2.**
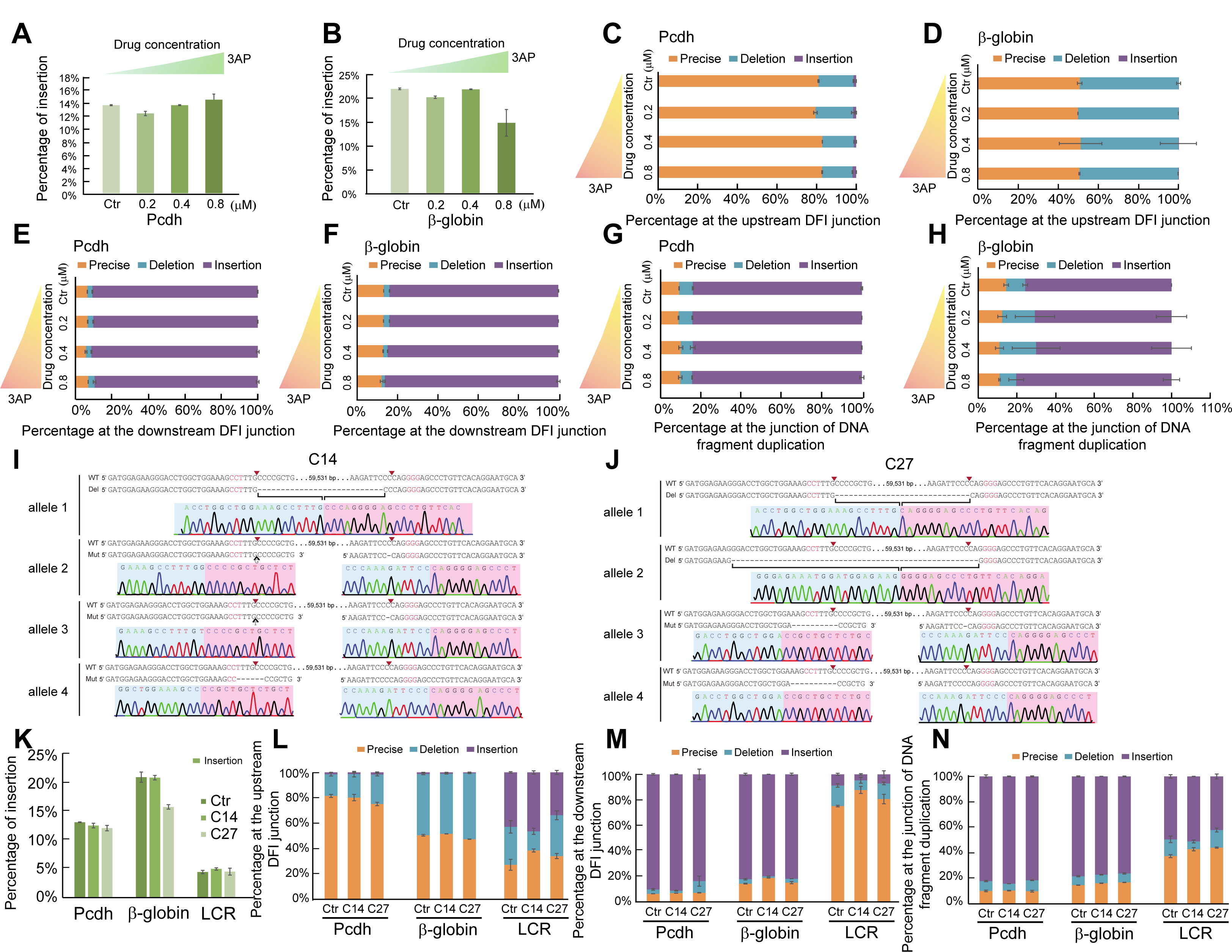
Analyses of DFE Junctions with 3AP Inhibition of CtIP Activity and in CtIP Mutant Cell Clones, Related to Figure 2. (A-B) 3AP results in no significant alteration of insertion at the DFD junctions. (C-H) 3AP does not alter the precise ligations and indels at the DFI junctions as well as DNA fragment duplication junctions. (I-J) Generation of subcloned cell lines of the CtIP mutations using the CRISPR/Cas9 system with a pair of sgRNAs. Shown are Sanger sequencing traces for two CtIP mutant CRISPR cell clones (C14 and C27). Sequencing confirmed the CtIP mutations. (K-N) There is no significant alteration of insertions at DFD junctions as well as of precise ligations and indels at the DFI junctions and DNA-fragment-duplication junctions in CtIP mutant cell clones. Data as mean ± SD.

**Figure S3.**
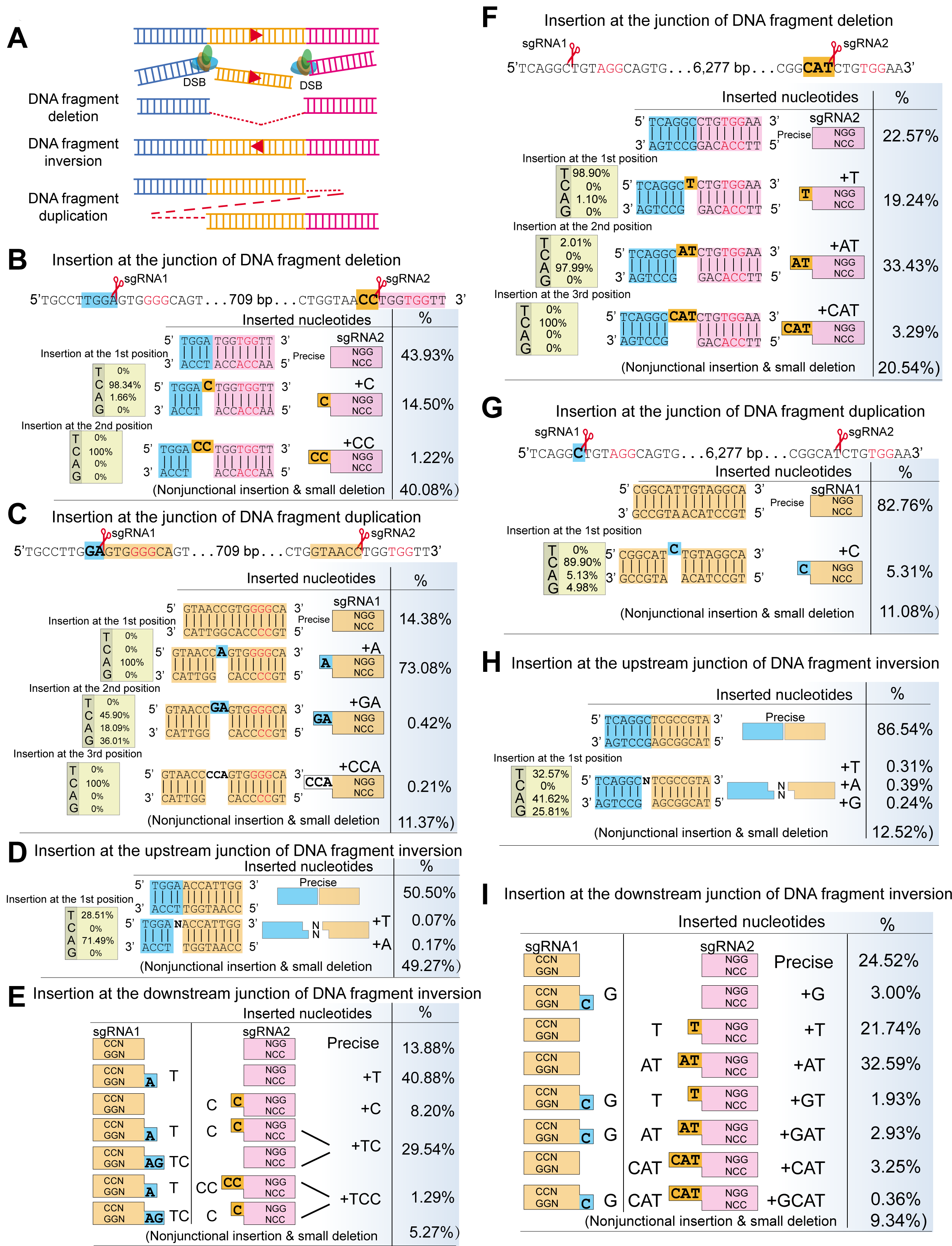
Analyses of Nucleotide Insertions at the Junctions of DFE of 709-bp or 6,277-bp DNA Fragments in the Human *β-globin* Locus, Related to Figure 3. (A) Shown are the schematics of the DNA-fragment deletion, inversion, and duplication. (B) Analyses of the insertions at DFD junctions of a 709-bp DNA fragment. The inserted nucleotides at DFD junctions are resulted from the cleavages by RuvC of the noncomplementary strand upstream of the **second** PAM site. The sequences of a targeted *β-globin* DNA fragment are shown with PAM in red. The inserted nucleotides detected by NGS are highlighted in the yellow background and are shown with their frequencies. The relative ratio of “T”, “C”, “A”, “G” at each position of insertions is also shown. (C) Analyses of insertions at the junctions of a 709-bp DNA fragment duplication. The inserted nucleotides at the junctions of DNA fragment duplications are resulted from the cleavages by RuvC of the noncomplementary strand upstream of the **first** PAM site. The inserted nucleotides are highlighted in the blue background and are shown with their frequencies. (D) Nonspecific insertions at the upstream junctions of DFI of a 709-bp DNA fragment. (E) The inserted nucleotides at the downstream junctions of DFI of a 709-bp DNA fragment are shown with their frequencies. They are resulted from the combined insertions at junctions of both DNA-fragment deletion (Cas9 with sgRNA2) and duplication (Cas9 with sgRNA1). (F-I) Analyses DFE junctions of a 6,277-bp DNA fragment reveal similar principles of Cas9-mediated nucleotide insertions.

**Figure S4.**
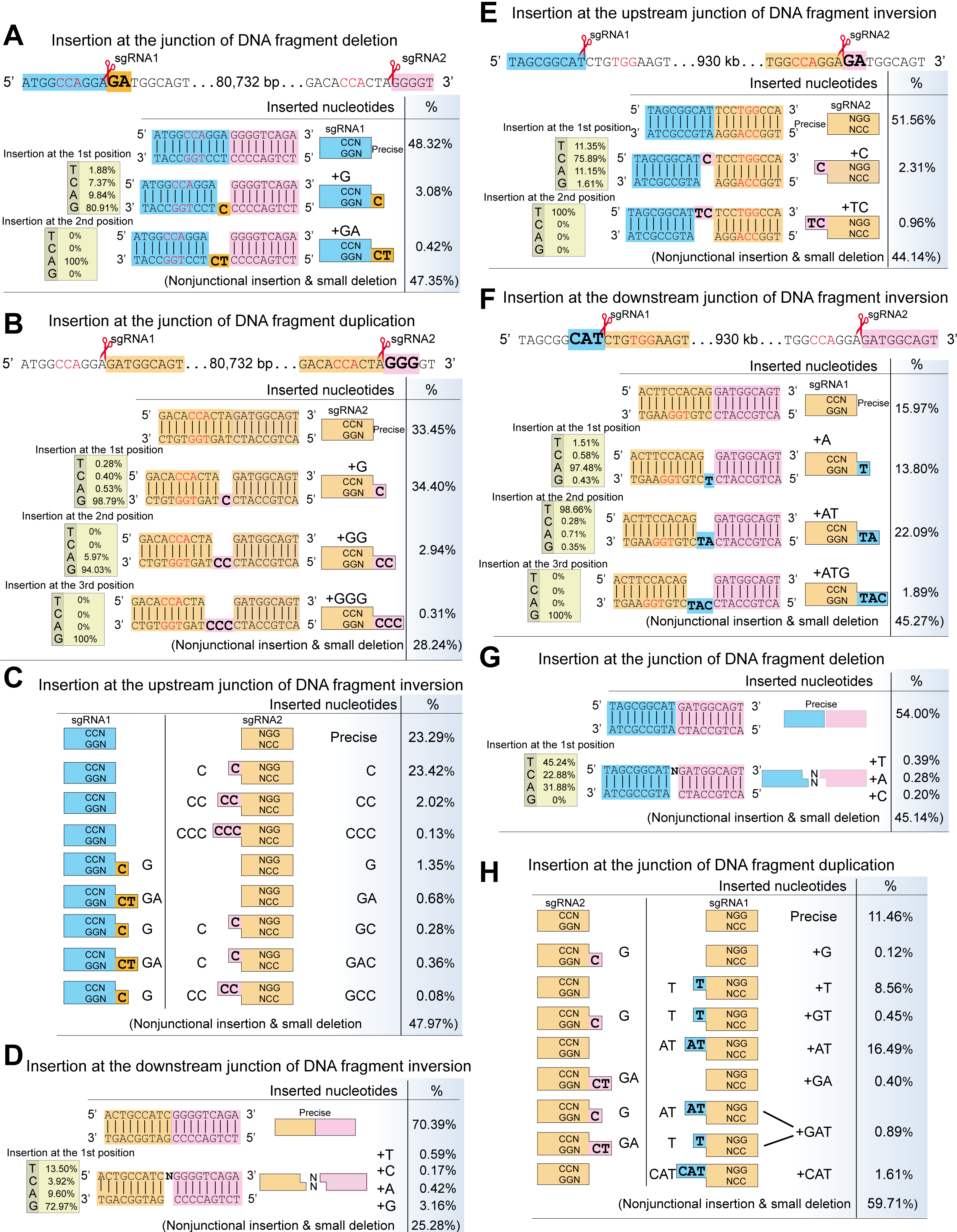
Analyses of Nucleotide Insertions at the Junctions of DFE of 80,732-bp or 930-kb DNA Fragments with Distinct PAM Configurations in the Human *β-globin* Locus, Related to Figure 3. (A) Analyses of insertions at the DFD junctions of an 80,732-bp DNA fragment. The inserted nucleotides are resulted from the cleavages by RuvC of the noncomplementary strand upstream of the **first** PAM site. The inserted nucleotides are highlighted in the yellow background and are shown with their frequencies. The relative ratio of “T”, “C”, “A”, “G” at each position of insertions is also shown. (B) Analyses of insertions at the junctions of an 80,732-bp DNA-fragment duplication. The inserted nucleotides are resulted from the cleavages by RuvC of the noncomplementary strand upstream of the **second** PAM site. The inserted nucleotides are highlighted in the pink background and are shown with their frequencies. (C) Analyses of insertions at the **upstream** junctions of DFI of an 80,732-bp DNA fragment. The inserted nucleotides are shown with their frequencies. They are resulted from the combined insertions at junctions of both DNA-fragment deletion (Cas9 with sgRNA1) and duplication (Cas9 with sgRNA2). (D) Nonspecific insertions at the downstream junctions of DFI of an 80,732-bp DNA fragment. (E) Analyses of nucleotide insertions at the upstream junctions of a 930-kb DNA fragment inversion. The inserted nucleotides are resulted from the cleavages by RuvC of the noncomplementary strand upstream of the **second** PAM site. (F) Analyses of nucleotide insertions at the downstream junctions of DFI of a 930-kb DNA fragment. The inserted nucleotides are from the cleavages by RuvC of the noncomplementary strand upstream of the **first** PAM site. The inserted nucleotides are highlighted in the blue background and are shown with their frequencies. (G) Nonspecific insertions at the junctions of DFD of a 930-kb DNA fragment. (H) Analyses of insertions at the junctions of DNA fragment **duplication of** a 930-kb DNA fragment. The inserted nucleotides are derived from the combined insertions at the both upstream (Cas9 with sgRNA2) and downstream junctions of DFI (Cas9 with sgRNA1).

**Figure S5.**
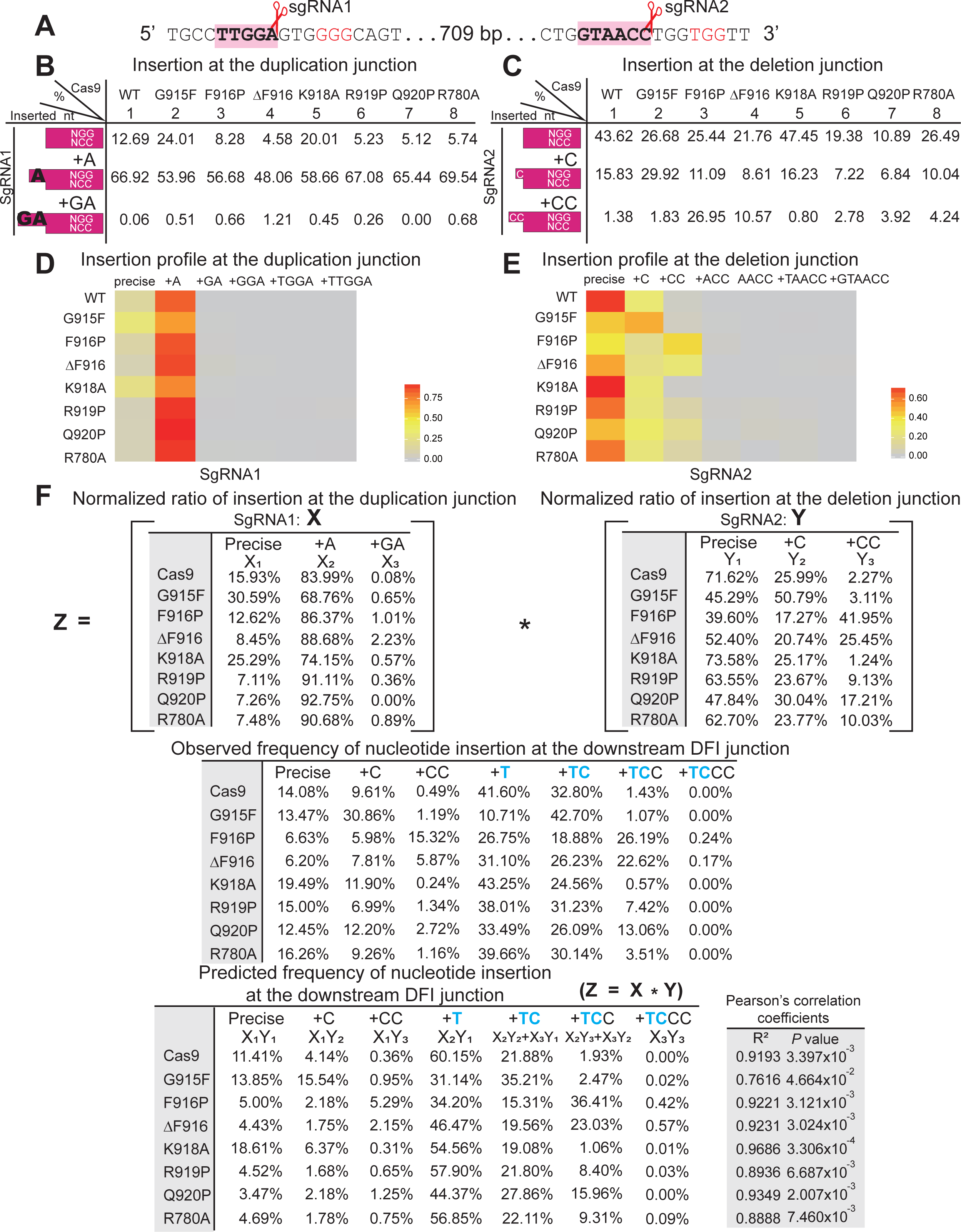
Altered Scissile Profiles of Rationally Engineered Cas9 Nucleases Reveal Predictable DFEs of a 709-bp Fragment in the Human *β-globin* Locus, Related to Figure 5. (A) The sequences of a targeted DNA fragment are shown with PAM in red. (B-C) Shown are the inserted nucleotides at the duplication or deletion junctions and their frequencies. (D-E) The heatmap of the relative ratio of insertions at the duplication and deletion junctions reveals altered scissile profiles of engineered Cas9 nucleases. (F) Predictable insertions at the downstream DFI junctions. The observed frequencies of the downstream insertions at the DFI junction are predicted to be equal to the product of multiplication of the normalized insertion frequencies at the both duplication and deletion junctions. Note that the insertions of “+TC” and “+TCC” are each from two different combinations. The Pearson’s correlation coefficients and their *P* values for all of the engineered Cas9 nucleases are indicated on the right.

**Figure S6.**
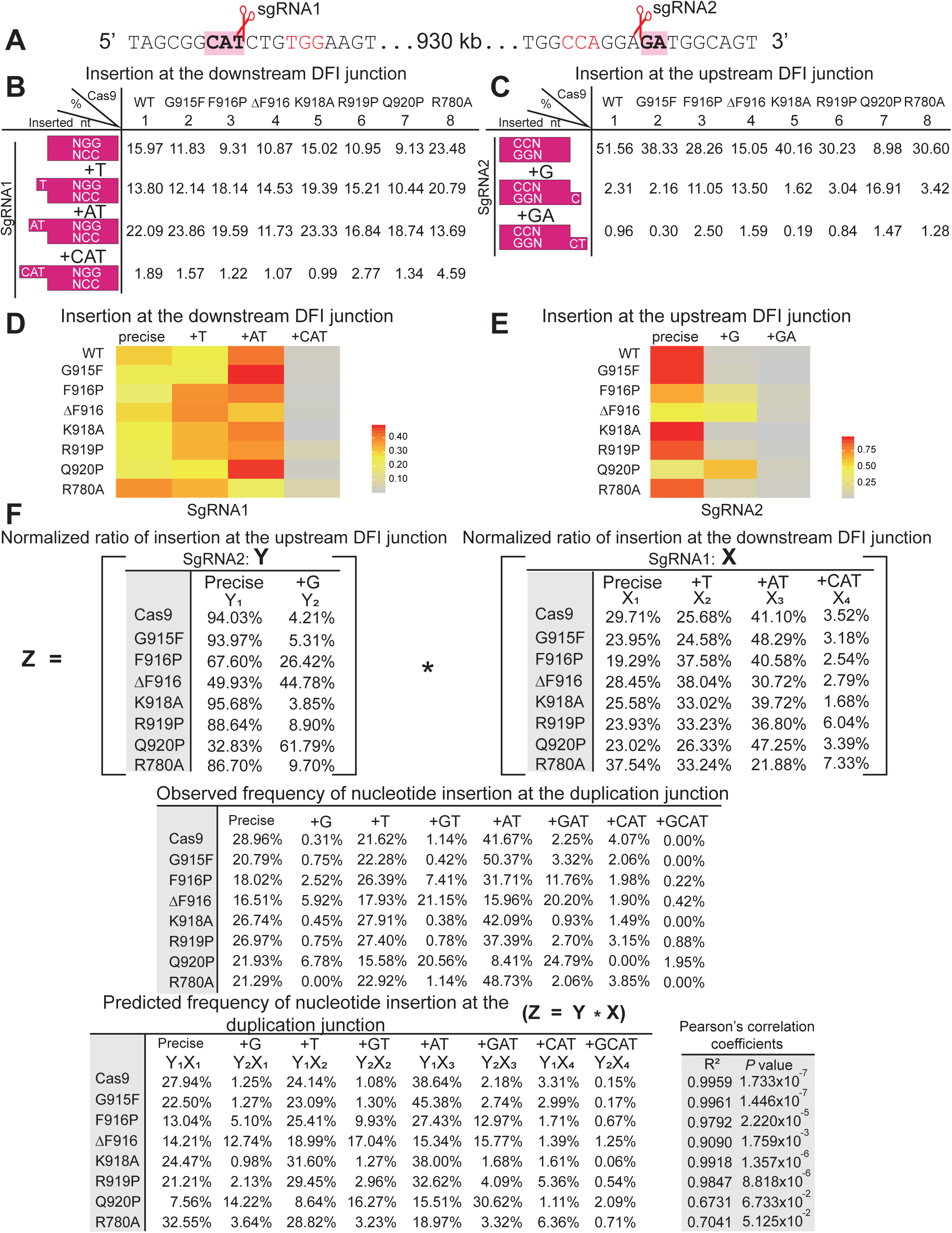
Altered Scissile Profiles of Rationally Engineered Cas9 Nucleases Reveal Predictable DFEs of a 930-kb Fragment in the Human *β-globin* Locus, Related to Figure 5. (A) The sequences of a targeted *β-globin* DNA fragment are shown with PAM in red. (B-C) Shown are the inserted nucleotides at the downstream and upstream junctions of DFI as well as their frequencies. (D-E) The heatmap of the relative ratio of insertions at the downstream and upstream junctions of DFI reveals altered scissile profiles of engineered Cas9 nucleases. (F) Predictable insertions at the **duplication** junctions. The observed frequencies of the insertions at the duplication junctions are predicted to be equal to the product of multiplication of the normalized insertion frequencies at the both upstream and downstream junctions of DFI. The Pearson’s correlation coefficients and their *P* values for all of engineered Cas9 nucleases are indicated on the right.

**Figure S7.**
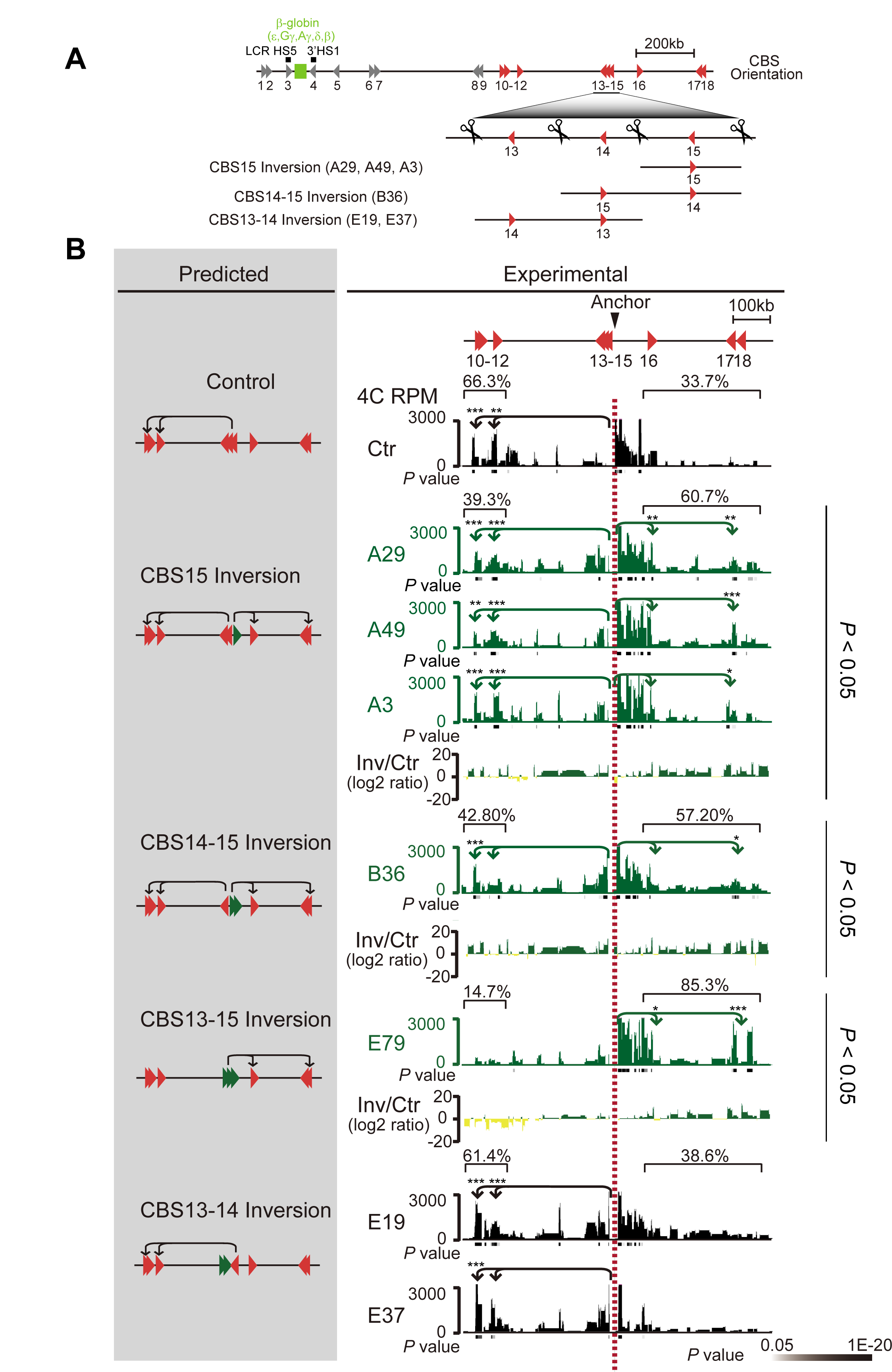
DFI of Progressive Numbers of Tandem CBS Sites at the Border Region of a *β-globin* Topological Domain Reveals Principles of CBS Insulation, Related to Figure 6. (A) Schematic of the orientated CBS clusters and the various DFIs of CBS combinations containing the inversion of CBS15, CBS14-15, or CBS13-14 induced by paired sgRNAs in the *β-globin* locus. (B) Shown are the schematics of predicted long-distance chromatin-looping interactions by inverting the relative orientation of CBSs in the *β-globin* locus. The chromatin-interaction profiles in control (Ctr) and various CBSs inversion clones using CBS13-15 as anchor are shown in the right panel. The CBS13-15 cell clone (E79) is a positive control (Guo et al., 2015). Note that the inversion of the two internal CTCF sites of CBS13-14 does not switch the chromatin-looping direction. Only inversions covering the boundary CBS15 switch the chromatin-looping direction. The significance of interactions (*P* value) is shown under the reads density. The log2 ratios between each inversion and the control clone are also indicated. \**P* < 0.05, \*\**P* < 0.01, \*\*\**P* < 0.001.

**Supplemental Table S1.** Oligonucleotides Used in This Study, Related to Experimental Procedures

